# YaHS: yet another Hi-C scaffolding tool

**DOI:** 10.1101/2022.06.09.495093

**Authors:** Chenxi Zhou, Shane A. McCarthy, Richard Durbin

**Affiliations:** Department of Genetics, University of Cambridge, Downing Street, Cambridge, CB2 3EH, UK; Wellcome Sanger Institute, Wellcome Genome Campus, Hinxton, Cambridge, CB10 1SA, UK

## Abstract

We present YaHS, a user-friendly command-line tool for construction of chromosome-scale scaffolds from Hi-C data. It can be run with a single-line command, requires minimal input from users (an assembly file and an alignment file) which is compatible with similar tools, and provides assembly results in multiple formats, thereby enabling rapid, robust and scalable construction of high-quality genome assemblies with high accuracy and contiguity. YaHS is implemented in C and licensed under the MIT License. The source code, documentation and tutorial are available at https://github.com/c-zhou/yahs.

## 1 Introduction

The rapid revolution of long-read, single-molecule DNA sequencing technologies in read length, base accuracy and per base cost is driving a golden age for *de novo* genome assembly. Multiple genome sequencing projects have been launched in the past few years, such as the Earth Biogenome Project (EBP, Lewin *et al*., 2018), the Vertebrate Genomes Project (VGP, Rhie *et al*., 2021), and the Darwin Tree of Life Project (DToL, Blaxter *et al*., 2022) aiming to assemble high quality, chromosome-scale genomes for many thousands of species across a range of genome sizes, complexity and ploidy. Despite the technological advances, assembly of reference quality genomes with long-read sequencing data alone remains elusive (Amarasinghe *et al*., 2020). Long-distance linkage information, such as physical maps, genetic maps, optical maps and Hi-C contact maps, is often used to construct chromosome-scale scaffolds from contigs. Hi-C is a sequencing-based proximity ligation assay that provides contact information between pairs of loci, originally designed to study the 3D structure of the genome inside a cell nucleus (Lieberman-Aiden *et al*., 2009). Since the contact frequency between loci pairs strongly correlates with separation on the genome, Hi-C has rapidly gained popularity as an economical method for generating chromosome-scale scaffolds (Burton *et al*., 2013; Dudchenko *et al*., 2017). Several scaffolding tools have been developed for construction of chromosome-scale assembly with Hi-C data including LACHESIS (Burton *et al*., 2013), HiRise (Putnam *et al*., 2016), 3D-DNA (Dudchenko *et al*., 2017), SALSA2 (Ghurye *et al*., 2019), and pin hic (Guan *et al*., 2021). Each of these has its own limitations and the results are affected by various factors such as genome complexity and repeat content, Hi-C library preparation and sequencing coverage (Ghurye *et al*., 2019; Kadota *et al*., 2020; Guan *et al*., 2021).

In this article, we introduce YaHS, another scaffolding tool which constructs chromosome-scale scaffolds utilising Hi-C data. YaHS follows a standard framework of Hi-C scaffolding pipelines: map Hi-C reads to input contigs, break contigs where necessary to correct assembly errors, build a contact matrix, construct and prune a scaffolding graph and finally output scaffolds. The core idea behind YaHS, which distinguishes it from other Hi-C scaffolding tools, is a novel method for building the contact matrix (Supplementary Methods). This method enables more accurate inferences of contig joins. In comparisons with previous tools applied to both simulated and real data, YaHS generated genome assemblies of higher accuracy and contiguity, and was more robust to assembly errors.

## 2 Results

### 2.1 Overview

The scaffolding process starts with mapping Hi-C reads to the input contigs, which falls outside of the scope of YaHS. The Arima Genomics’ mapping pipeline was employed in this study which consists of four major steps: read mapping in single-end read mode, read filtering for chimeric joins across ligation junctions, read pairing and PCR duplicate removal (https://github.com/ArimaGenomics/mapping_pipeline). YaHS takes the alignment file (either in BED format or BAM format) to first optionally break contigs at positions lacking Hi-C coverage which are potential assembly errors. Scaffolding then proceeds in multiple rounds. In each round YaHS builds a contact matrix by splitting each contig into chunks of a certain size (i.e., resolution) and assigns Hi-C contact signals into cells of chunk pairs. Here we refer to cells within contigs as intra-cells and between contigs as *inter*-cells. The Hi-C contact frequencies are counted for each cell. To calculate the joining score of a pair of contigs, the contact frequencies of the *inter*-cells between them are normalised by expected values which are estimated by the medians of the intra-cells at the same separations and then used to calculate a weighted sum. The fundamental idea is that the *inter*-cells of neighbouring contigs on a scaffold should have similar contact frequencies to intra-cells at the same separations. YaHS also optionally takes account of the restriction enzymes used in the Hi-C library, and if so, the cell contact frequencies are normalised first by the corresponding number of cutting sites. YaHS next builds a scaffolding graph with contigs as nodes and contig joins as edges which are weighted by the joining scores calculated in the previous step. The graph is simplified by a series of operations including filtering low score edges, trimming tips, trimming blunt ends, solving repeats, removing transitive edges, removing bubbles, resolving ambiguous orientations, trimming weak edges and removing ambiguous edges. Finally the graph is traversed to assemble scaffolds along contiguous paths. A second step of assembly error correction is optionally performed to break scaffolds at positions of contig joins without sufficient Hi-C coverage. YaHS runs multiple rounds of scaffolding at decreasing resolution (increasing chunk sizes). See Supplementary Methods for further details.

### 2.2 On simulated human genome assemblies

We randomly split the Telomere-to-Telomere (T2T) human genome assembly (T2T-CHM13, Nurk *et al*., 2022) into 100Kb to 1Mb chunks and ended up with a simulated assembly of 5,483 contigs with an N50 of 715Kb. The T2T Arima Hi-C data was downloaded from NCBI (accession SRX10230901-SRX10230903) and mapped to the simulated assembly for reconstruction of the genome with YaHS, SALSA2 and pin hic. YaHS assembled over 92% sequences into 25 major scaffolds (¿35Mb). The N50 and N90 were 132.6Mb (L50 = 9) and 36.5Mb (L90 = 23) respectively (Fig. 1A). In contrast, the N50 and N90 of SALSA2 assembly were 43.9Mb (L50 = 22) and 3.3Mb (L90 = 103) respectively (Fig. 1B), and the N50 and N90 of pin hic assembly were 51.5Mb (L50 = 19) and 8.5Mb (L90 = 68) respectively (Fig. 1C). We used QUAST-LG (Mikheenko *et al*., 2018) to map the scaffolds to the T2T-CHM13 genome assembly for quality assessment. Three types of misassemblies were considered: relocations (gaps on scaffolds), inversions (misorientated contigs) and translocations (misjoins of inter-chromosomal contigs). The numbers of relocations and inversions reported by QUAST-LG for YaHS, SALSA2 and pin hic assemblies were 40 and 21, 262 and 55, 171 and 115, respectively. No translocation was reported for any of the three assemblies.

**Figure 1.**
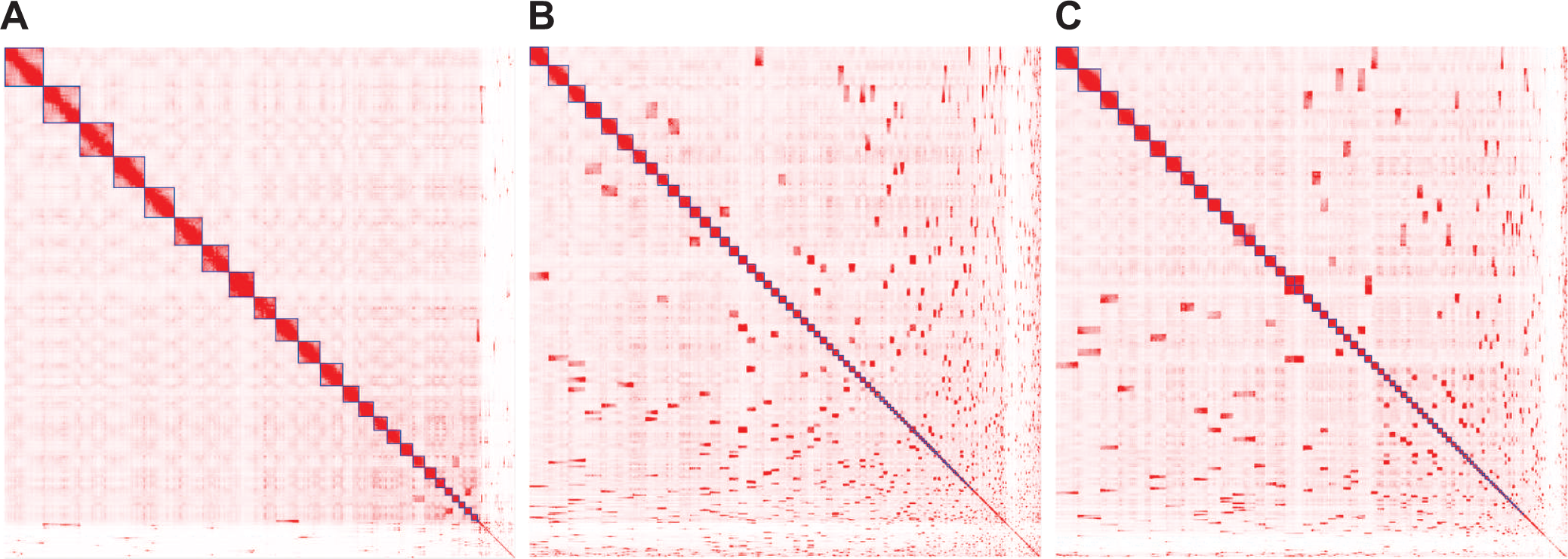
Hi-C contact maps of genome assemblies constructed with YaHS (A), SALSA2 (B) and pin hic (C) for the simulated T2T data without contig errors. The blocks highlighted with blue squares in diagonal line are scaffolds. The contact maps were plotted with Juicebox (Durand *et al*., 2016).

To evaluate the performance of these tools on assemblies with errors, we generated another assembly by randomly introducing 25 erroneous contigs into the previous one. These included ten with a single misjoin of two *intra*-chromosomal contigs, ten with a single misjoin of two *inter*-chromosomal contigs and five with two misjoins of any three contigs - a total of 30 assembly errors. The joining orientations were randomly determined. In total these affected 32.1Mb of contig sequences. The new assembly comprised 5,453 contigs with an N50 of 718Kb. YaHS corrected 28 errors out of the 30. The two errors missed were on a double-misjoined contig with two closely located contigs from chr10 (a gap of 1.2Mb) flanking a short contig (170Kb) from chr19. The error detection in this scenario was more challenging. SALSA2 corrected 14 errors out of the 30. Pin hic did not do assembly error correction explicitly but instead broke scaffolds at suspicious misassembled positions at the end of the scaffolding process. The contiguity of the scaffolding resulting from each the three tools was similar to that of the previous error-free assembly: all statistics for YaHS remained identical; SALSA2 had a slightly larger N50 of 46.1Mb and smaller L50 of 20; pin hic ended up with a more fragmented assembly with N50 and N90 respectively decreasing to 37.5Mb (L50 = 28) and 5.8Mb (L90 = 98). The scaffolding errors for YaHS reported by QUAST-LG also remained similar with two more relocations and two extra translocations due to the uncorrected contig errors. In contrast, significantly more errors were reported for the other two assemblies. The numbers of relocations, inversions and translocations were 278, 18 and 63 respectively for the SALSA2 assembly, and 188, 22, and 123 respectively for the pin hic assembly.

### 2.3 On Darwin Tree of Life assemblies

We applied the three tools to the construction of genome assemblies for 15 DToL species across a range of taxonomic groups, genome sizes and initial assembly quality. YaHS consistently generated assemblies with higher contiguity compared to SALSA2 and pin_hic particularly for the L90 statistics (Supplementary Table S1, Supplementary Figures S1-S15).

In particular, we constructed a genome assembly for the oak bush-cricket (*Meconema thalassinum*, ToLID: iqMecThal1) which has a very large genome size estimated to be over 9Gb. The initial genome assembly consisted of 2,093 contigs of 9,054Mb with a N50 of 10.7Mb (L50 = 229). YaHS detected 268 assembly errors. In the final assembly, three scaffolds longer than 1,349 Mb constituted over 50% of the assembly, and 13 scaffolds longer than 179.6 Mb constituted 90% of the assembly. The largest scaffold was 2,087Mb. In comparison, the N50 and N90 for the SALSA2 assembly were 79.0Mb (L50 = 18) and 3.5Mb (L90 = 311) respectively, and for the pin hic assembly were 202.4Mb (L50 = 12) and 14.2Mb (L90 = 82) respectively. See Supplementary Figure S1 for the Hi-C contact maps of these genome assemblies.

## 3 Conclusion

YaHS is a fast, reliable and accurate tool for construction of chromosome-scale scaffolds with Hi-C data that is now being used routinely by the DToL project and others. It consistently outperforms other state-of-the-art Hi-C scaffolding tools in both genome assembly accuracy and contiguity across a wide range of species and genome sizes, and initial assembly quality. It learns its parameters from the data so is robust to Hi-C data with different genomic separation distributions, including generated with different protocols. It is open source, easy to use and well-documented.

## Acknowledgements

The authors thank Marcela Uliano-Silva, Ksenia Krasheninnikova, João Gabriel, Jonathan Wood, Alan Tracey, Camilla El-dridge, Bethan Manley and Martin Pippel, who used the first versions of the tool and provided much valuable feedback to help improve the usability.

