## Supplemental materials for "YaHS: yet another Hi-C scaffolding tool"

#### Supplementary materials

### 1 Supplementary methods

#### 1.1 Assembly error correction

To detect potential assembly errors for a contig, we first count the Hi-C read pairs spanning each position. Instead of using all read pairs, we require the mapping distance of two reads to fall under a certain threshold in order to reduce the influence of background noise. The distance threshold is estimated from data, such that 80% of read pairs are included. If this estimate is smaller than 1Mb, we use 1Mb instead. For each contig we then calculate the median value of the number of read pair counts crossing each site, and break the contig at positions where counts drop below a tenth of the median (except at the ends of the contig). The new contigs are subject to further rounds of error corrections until no errors are detected. See Fig. M1A.

#### 1.2 Contig joining scores

##### 1.2.1 Notation

Consider a genome assembly of $N$ contigs. Denote by $a_n$ the size of $n$th ($n = 1, ..., N$) contig. For a predefined resolution $l$, each contig $n$ is split into $b_n$ chunks of size $l$, where $b_n = \lceil \frac{a_n}{l} \rceil$. The Hi-C read pair count between chunks $i$ and $j$ is denoted by $c_{i,j}$, and the distance of separation of chunk $i$ to $j$ by $d_{i,j} = \delta(i,j) = |i - j|$, where $i, j \in 1, ..., b_n$. The definition of $c_{i,j}$ and $d_{i,j}$ can be easily extended to a pair of contigs $m$ and $n$, e.g., $\delta(i,j) = b_m - i + j$ when $m$ is followed by $n$ in their positive orientations, and we have $i \in 1, ..., b_m$ and $j \in 1, ..., b_n$. For convenience, a chunk pair $(i,j)$ is referred to as a cell, and a cell for a chunk pair within a contig or between two contigs is respectively referred to as an *intra*- or *inter*-cell. We should note that, the subscripts for contigs are dropped here for convenience, e.g., the *intra*- and *inter*-cell Hi-C count should be $c_{n,i,j}$ and $c_{m,n,i,j}$, respectively. See Fig. M1B.

##### 1.2.2 The expected Hi-C count of a cell

The expected Hi-C count of a cell $(i,j)$ is considered a function of chunk distance $d_{i,j}$. The cells with the same distances form a band in the Hi-C contact map, which we assume have the same expected values denoted by $E_d$. We use *intra*-cells to estimate this value $E_d = Med\{c_{n,i,j} : n \in 1, ..., N, \delta(i,j) = d\}$, where the function $Med$ calculates the median value of a data set. We also explored using the average values as expectations. For contig $n$, the total Hi-C count of band $d$ is written as $C_{n,d} = \sum_{i,j:\delta(i,j)=d} c_{i,j}$, and the total cell count of band $d$ is written as $B_{n,d} = \sum_{i,j:\delta(i,j)=d} 1 = b_n - d$, where $i, j \in 1, ..., b_n$. Then the expected Hi-C count of cell $(i,j)$ with $\delta(i,j) = d$ is averaged across all contigs and calculated as $E_d = \sum_{n=1}^{N} C_{n,d} / \sum_{n=1}^{N} B_{n,d}$. We found that the average values performed less well compared to the medians for using as expectations.

##### 1.2.3 The score for contig joining

We consider the joining of a contig pair $m$ and $n$ in four possible orientations and the one with the largest score is selected as the final orientation. Without loss of generality, in this subsection, we assume the joining of the negative strand of $m$ followed by the positive strand of $n$. The joining score is calculated as the weighted sum of the normalised Hi-C counts of the *inter*-cells between the two contigs. The Hi-C count of *inter*-cell $(i,j)$ (chunk $i$ from contig $m$ and chunk $j$ from contig $n$), where $i \in 1, ..., \lfloor \frac{b_m}{2} \rfloor$ and $j \in 1, ..., \lfloor \frac{b_n}{2} \rfloor$, is normalised by the expected value and written as $w_{i,j} = c_{i,j} / E_{\delta(i,j)}$. Similar to the *intra*-cell calculation, the total cell count of band $d$ is denoted by $B_{m,n,d} = \sum_{i,j:\delta(i,j)=d} 1$. Intuitively, we want the Hi-C signals to fill the Hi-C contact map band to band from inner to outer, and therefore define a filling rate up to band $d$ as $f_{m,n,d} = \sum_{i,j:\delta(i,j)<d} w_{i,j} / \sum_{k=1}^{d-1} B_{m,n,k}$. The joining score of contig $m$ and $n$ is finally calculated as $s_{m,n} = \sum_{i,j:\delta(i,j)<D} f_{m,n,\delta(i,j)} w_{i,j} / \sum_{k=1}^{D} B_{m,n,k}$, where $D = \min(\text{argmax}_d(\sum_u B_{u,d} \geq 30), \text{argmin}_d(\sum_u \sum_{v=1}^{d} C_{u,v} / \sum_u \sum_v C_{u,v}) \geq 0.99)$, is the distance threshold calculated from data - either the maximum value of $d$ such that the total number of *intra*-cells on band $d$ is no less than 30 or the minimum value of $d$ such that the number of Hi-C read pairs fall into bands with distance no greater than $d$ constitute at least 99% of the total number of Hi-C read pairs, whichever is smaller.

##### 1.2.4 Remove background noise

Background noise is considered random Hi-C contacts between loci which could be either due to library preparation or read mapping issues. We see noise levels vary significantly between different species and Hi-C data types. We estimate a global value for noise level from the data using *inter*-cells during the joining score calculation. More precisely, after finishing Hi-C counting for each pair of contigs, we choose the orientation with the least Hi-C counts from the four possibilities and record the counts for Hi-C contacts and cells. The average of cell Hi-C count across all contig pairs (most of which will be on different chromosomes) is used as the estimation for

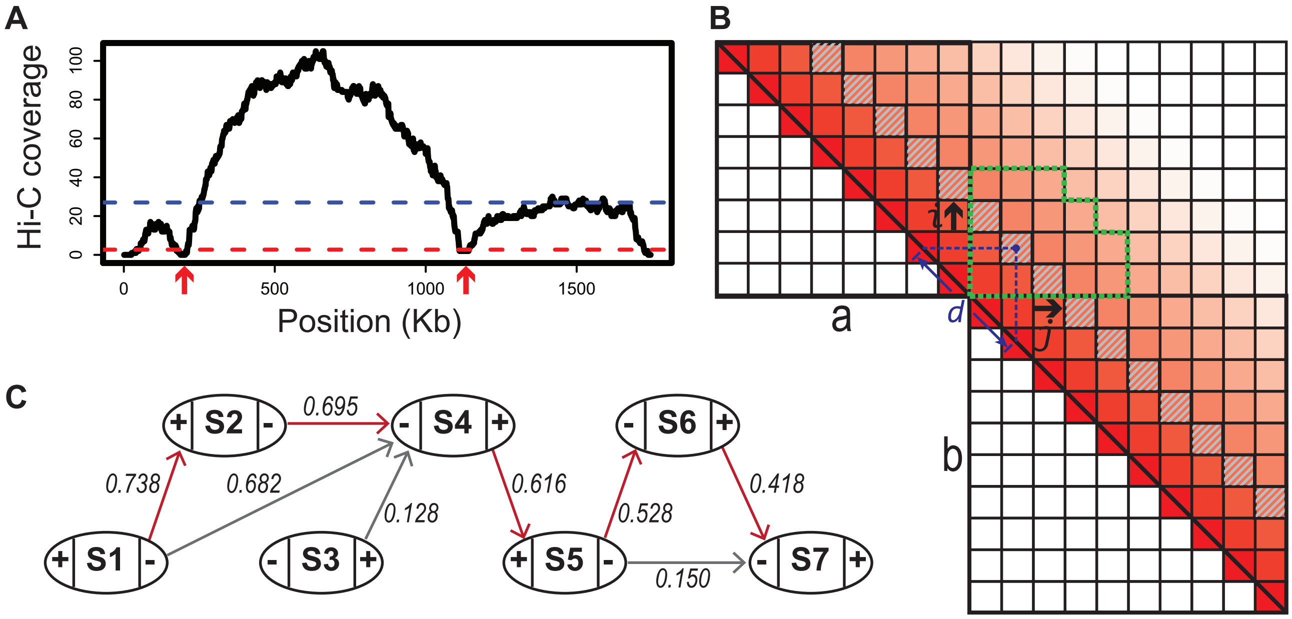


Figure M1: An overview of YaHS. (**A**) Number of Hi-C read pairs spanning the positions along a contig. The horizontal dashed lines indicate the median value of coverage (blue) and the lower threshold for detecting assembly errors which is a tenth of the median value (red). The red arrows indicate positions of contig breaks. (**B**) Contigs **a** and **b** are partitioned into eight and ten chunks respectively, and the Hi-C signals are consequently assigned to 36 and 55 *intra*-cells and 80 *inter*-cells (only consider the upper triangular part). The color of a cell indicates the Hi-C contact frequency (darker colors for higher frequencies). In general the contact frequency decreases as the distance of separation ($d$) between chunks increases. If **a** and **b** are neighboring contigs on a scaffold then cells with the same separation, e.g. those cells shaded with gray slashes ($d = 3$), should have similar contact frequencies whether they are *intra*- or *inter*-cells. Specifically, *inter*-cells within the green rectangle are used to calculate the joining score of contig **a** and **b** in this orientation. Here the separation threshold for contig joining score calculation is set as 6, which in the program is estimated from the data. Scores for each of the four possible pairwise orientations are calculated and compared to determine the orientation of the join. (**C**) A scaffolding graph for seven contigs. Numbers on edges are joining scores. The path indicated by red edges is used to assemble a scaffold after graph simplification.

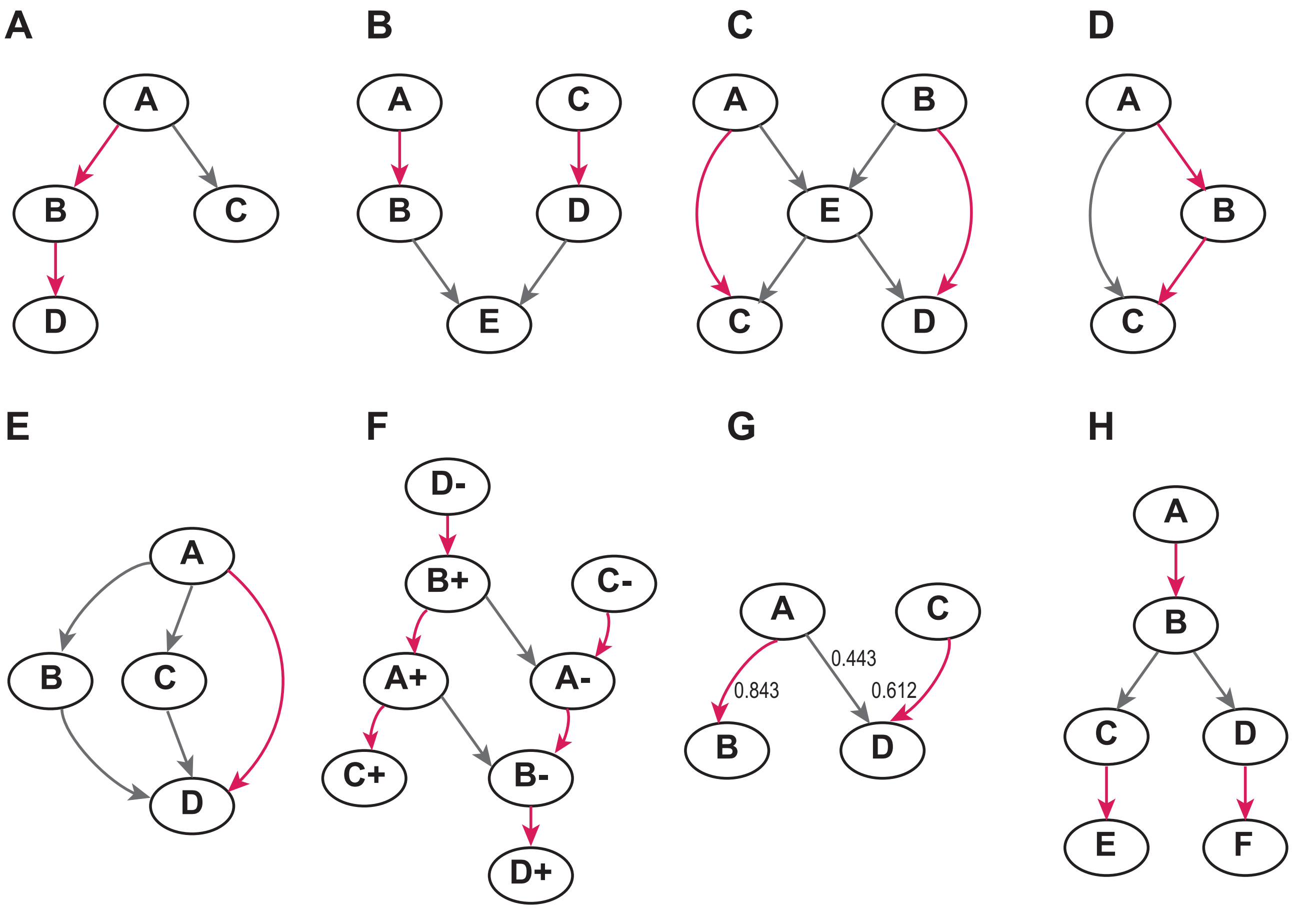


Figure M2: Examples for heuristic operations in scaffolding graph simplification for tips (A), blunt ends (B), repeats (C), transitive edges (D), bubbles (E), ambiguous orientations (F), weak edges (G), and ambiguous edges (F). The grey edges are removed after the operation in each example, leaving the red edges.

noise level, denoted by $e$. With the noise level included, the *inter*-cell Hi-C count normalisation is then written as $w_{i,j} = (c_{i,j} - e)/(E_{\delta(i,j)} - e)$.

##### 1.2.5 Consider Hi-C library restriction enzymes

The restriction enzymes used in the Hi-C library are optionally considered to remove biases on cell Hi-C counts caused by the variations in cutting site densities. We scan the input contigs to obtain the total number of cutting sites, denoted by $R$, and also to record the positions on contigs. The per base cutting site density is calculated as $r = R/G$, where $G$ is the total number of base pairs of the input contigs. For each chunk along a contig, we calculate the relative per base cutting site density as $r_{n,i} = \frac{R_{n,i}}{lr}$, where $R_{n,i}$ is the number of cutting sites in chunk $i$ of contig $n$. With the restriction enzymes considered, the cell Hi-C counts are normalised by the cutting site densities before any calculation mentioned above and written as $c'_{i,j} = c_{i,j}/r_i r_j$, dropping implied subscripts for contigs.

#### 1.3 Scaffolding graph construction and pruning

A scaffolding graph is constructed by using the contigs as nodes (two nodes for each contig for forward and reverse strands) and the joins between them as edges which are weighted by the joining scores (Fig. M1C). We implement nine heuristic operations for graph simplification.

(1) Filter by edge weight: edges with weights smaller than a predefined threshold or likely random contig joins are filtered out. To check if an edge is a likely random join, we find the 99th quantile, denoted by $q$, of a binomial distribution with $W$ trials and a probability of $w$ for success on each trial, where $W$ is the number of *inter*-cells between the two contigs and $w$ is the average of the normalised Hi-C counts for *inter*-cells across all contig pairs. The edge is considered a random join and filtered out if $pW < q$, where $p$ is the joining score.

(2) Trim edges for tips: a tip is a node with only one edge - either incoming or outgoing - and the edge connects to a node on a contiguous path. The edge connecting the tip is removed. See Fig. M2A.

(3) Trim edges for blunt ends: a blunt end is a node with only incoming or outgoing edges and each edge connects to a node on a contiguous path. Edges connecting the blunt end are all removed. See Fig. M2B.

(4) Solve repeats: a repeat is a chunk of sequence which occurs multiple times and can be placed in multiple locations. Reuse of repeats is error prone so we skip them and leave scaffolds with gaps. A repeat in the graph is identified as a node with multiple incoming and outgoing edges, where for each pair of incoming and outgoing nodes, there exists an edge connecting them. Edges connecting the repeat are all removed. See Fig. M2C.

(5) Remove transitive edges: suppose three contigs are placed on a scaffold in a linear order, then as well as there being Hi-C contacts between the first two contigs and between the last two contigs, it is also common to see Hi-C contacts between the fist and the third contig. In the graph, we call the edge connecting the first and the third node a transitive edge, which is removed. See Fig. M2D.

(6) Remove bubbles: a bubble structure in the graph involves a source and a sink node where there exists multiple paths between them and no incoming edges from or outgoing edges to other parts of the graph except at the source and the sink. It is not possible to decide which path to select. However, we can skip the bubble structure by directly joining the source and the sink and introduce a gap between them. See Fig. M2E.

(7) Resolve ambiguous orientations: an ambiguously orientated contig pair will end up with a diamond structure in the graph, e.g., the forward strand of the first contig points to both the forward and reverse strands of the second contig which points back to the reverse strand of the first contig (companion edges). Ambiguous edges are removed if the diamond structure can be resolved, e.g., if the first contig is the only incoming node for the forward strands of the second contig while there exists a third contig pointing to the reverse strand of the second contig, then the edge connecting the forward strand of first contig and the reverse strand of the second contig along with its companion edge are removed. See Fig. M2F.

(8) Trim weak edges: an edge is considered weak and removed if there exists at least another edge with a larger weight in both outgoing edges from the source node and incoming edges to the sink node. See Fig. M2G.

(9) Remove ambiguous edges: a set of at least two edges are considered ambiguous and removed if they share the same source or sink node. Once removed the ambiguous edges, all paths extracted from the graph traversal are unambiguous which are used to assemble scaffolds. See Fig. M2H.

#### 2 Supplementary tables

Table S1: Genome assembly results for the 15 DToL/VGP species. The cells of smaller L50 and L90 between YaHS and SALSA2 were coloured in green. Pin_hic was not taken into account for this colouring as it was sometimes too aggressive to make contig joins which led to obvious scaffolding errors (Fig. S1-S15).

| DToL/VGP ID | Accession | Taxa | Contig (Mb) | | | | | | YaHS scaffolds (Mb) | | | | SALSA2 scaffolds (Mb) | | | | Pin_hic scaffolds (Mb) | | | |
|---|---|---|---|---|---|---|---|---|---|---|---|---|---|---|---|---|---|---|---|---|
| | | | Size | NO. | N50 | L50 | N90 | L90 | N50 | L50 | N90 | L90 | N50 | L50 | N90 | L90 | N50 | L50 | N90 | L90 |
| iqMecThal1 | PRJEB48399 | Orthoptera | 9,054 | 2,093 | 10.7 | 229 | 2.4 | 925 | 1,349 | 3 | 179.6 | 13 | 79.0 | 18 | 3.5 | 311 | 202.4 | 12 | 14.2 | 82 |
| mMelMel3 | PRJEB49353 | Mammals | 2,662 | 614 | 75.2 | 12 | 8.1 | 47 | 118.7 | 9 | 44.2 | 22 | 110.0 | 9 | 8.9 | 38 | 124.9 | 5 | 35.3 | 18 |
| iqTelOcea10 | *pending* | Orthoptera | 2,018 | 7,906 | 0.6 | 887 | 0.1 | 4,126 | 120.0 | 6 | 27.2 | 16 | 16.4 | 24 | 0.4 | 365 | 13.5 | 39 | 1.3 | 205 |
| drGeuUrba1 | PRJEB48840 | Dicots | 1,347 | 493 | 20.4 | 24 | 5.7 | 70 | 64.8 | 9 | 45.0 | 19 | 65.2 | 9 | 45.0 | 19 | 65.9 | 4 | 43.2 | 15 |
| xgSteCine2 | PRJEB45667 | Molluscs | 1,303 | 885 | 6.2 | 57 | 1.0 | 256 | 68.5 | 9 | 34.1 | 18 | 31.7 | 13 | 1.8 | 81 | 63.5 | 8 | 5.3 | 26 |
| xgPhoLine1 | PRJEB48398 | Molluscs | 996.3 | 516 | 4.9 | 63 | 1.4 | 204 | 49.4 | 9 | 43.7 | 17 | 39.1 | 10 | 2.6 | 46 | 58.6 | 7 | 38.9 | 15 |
| wcLumRube1 | PRJEB53406 | Annelids | 914.8 | 2,678 | 0.7 | 376 | 0.2 | 1,352 | 43.1 | 8 | 0.6 | 71 | 11.0 | 21 | 0.5 | 211 | 4.0 | 60 | 0.7 | 257 |
| ihAcaHaem1 | PRJEB47321 | Hemiptera | 875.1 | 582 | 3.7 | 69 | 0.8 | 254 | 128.3 | 3 | 113.6 | 6 | 61.0 | 5 | 2.3 | 48 | 141.6 | 3 | 6.2 | 12 |
| icPodAlpi1 | PRJEB49177 | Coleoptera | 783.0 | 374 | 10.3 | 20 | 2.0 | 82 | 96.5 | 4 | 79.6 | 7 | 77.0 | 5 | 3.3 | 22 | 104.3 | 3 | 35.8 | 8 |
| idMelMell2 | PRJEB46300 | Diptera | 754.2 | 487 | 4.7 | 50 | 0.8 | 196 | 105.7 | 3 | 16.3 | 7 | 47.4 | 5 | 1.2 | 54 | 52.4 | 4 | 7.7 | 14 |
| drChaAngu1 | PRJEB47394 | Dicots | 663.8 | 175 | 26.1 | 10 | 13.7 | 23 | 38.2 | 7 | 24.3 | 15 | 21.7 | 12 | 10.7 | 29 | 39.9 | 4 | 32.2 | 11 |
| iyCerRyby1 | PRJEB45199 | Hymenoptera | 418.7 | 713 | 15.2 | 8 | 0.7 | 37 | 26.2 | 6 | 0.7 | 18 | 26.2 | 5 | 0.7 | 26 | 52.7 | 3 | 0.8 | 14 |
| ilPluXylo3 | PRJEB48401 | Lepidoptera | 323.7 | 38 | 11.0 | 14 | 6.1 | 28 | 11.3 | 13 | 8.6 | 26 | 11.1 | 13 | 6.6 | 27 | 11.7 | 7 | 8.6 | 20 |
| lpJunEffu1 | PRJEB50168 | Monocots | 243.9 | 271 | 2.4 | 32 | 0.6 | 107 | 11.6 | 10 | 9.6 | 19 | 5.9 | 13 | 0.9 | 51 | 13.4 | 4 | 8.5 | 14 |
| gfLaeSulp1 | PRJEB47319 | Fungi | 38.5 | 21 | 2.6 | 6 | 1.9 | 12 | 7.5 | 3 | 1.9 | 6 | 2.6 | 6 | 1.9 | 12 | 25.3 | 1 | 4.3 | 3 |

### 3 Supplementary figures

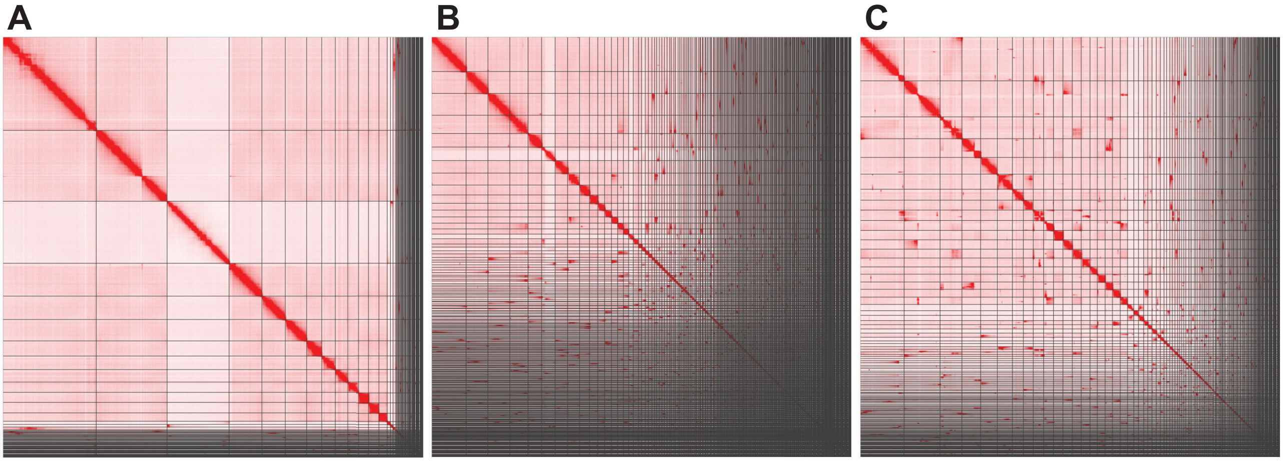


Figure S1: Hi-C contact maps of the oak bush-cricket (*Meconema thalassinum*, iqMecThal1) assemblies constructed from YaHS (A), SALSA2 (B) and pin_hic (C).

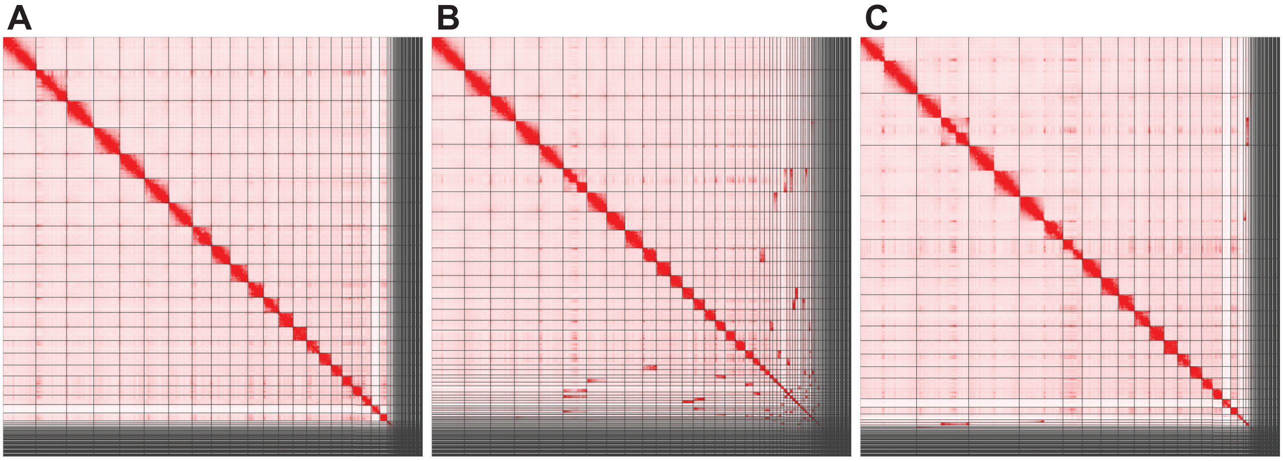


Figure S2: Hi-C contact maps of the European badger (*Meles meles*, mMelMel3) assemblies constructed from YaHS (A), SALSA2 (B) and pin_hic (C).

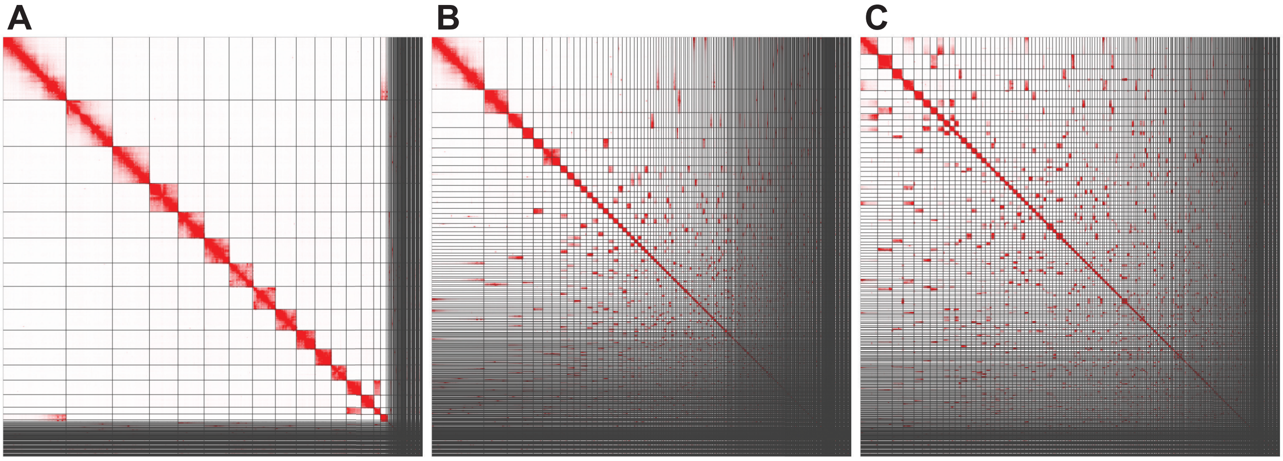


Figure S3: Hi-C contact maps of the black field cricket (*Teleogryllus oceanicus*, iqTelOcea1) assemblies constructed from YaHS (A), SALSA2 (B) and pin_hic (C).

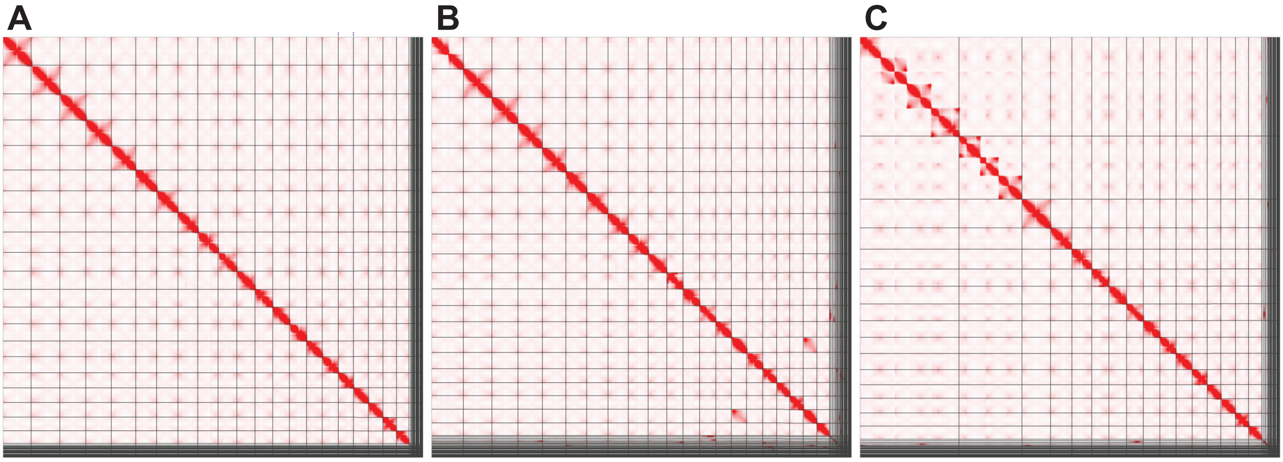


Figure S4: Hi-C contact maps of the wood avens (*Geum urbanum*, drGeuUrba1) assemblies constructed from YaHS (A), SALSA2 (B) and pin_hic (C).

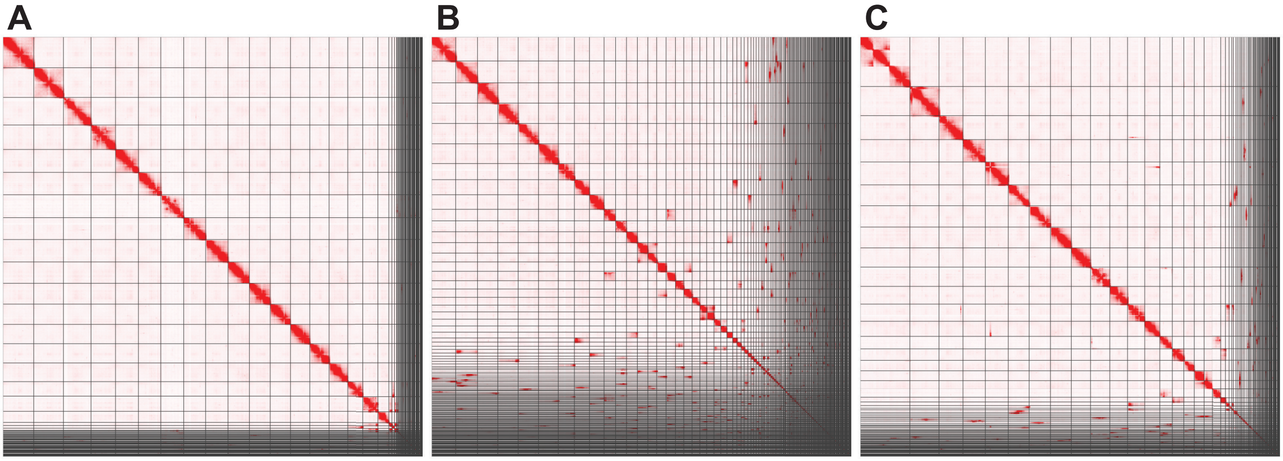


Figure S5: Hi-C contact maps of the grey top shell (*Steromphala cineraria*, xgSteCine2) assemblies constructed from YaHS (A), SALSA2 (B) and pin_hic (C).

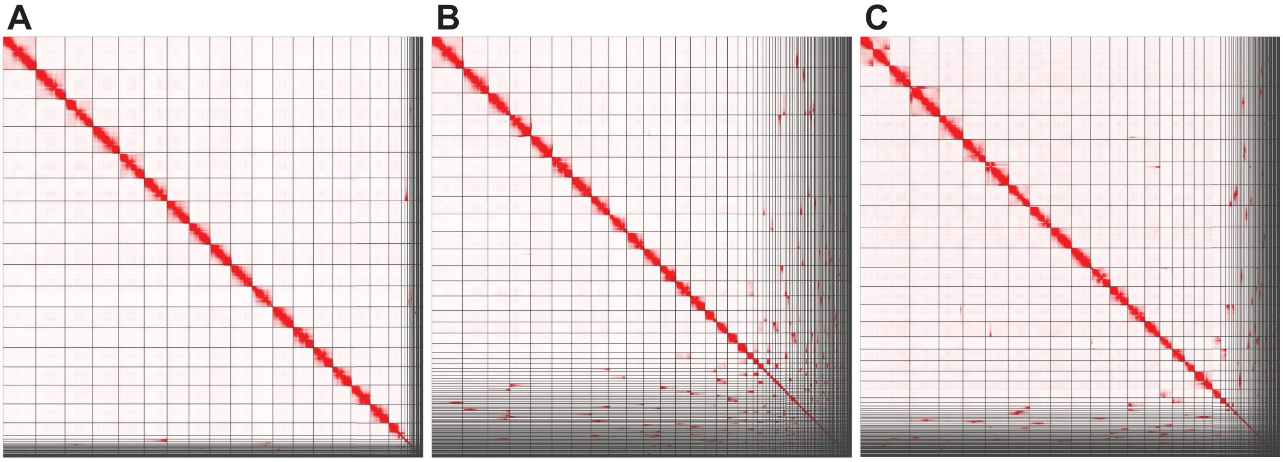


Figure S6: Hi-C contact maps of the thick top shell (*Phorcus lineatus*, xgPhoLine1) assemblies constructed from YaHS (A), SALSA2 (B) and pin_hic (C).

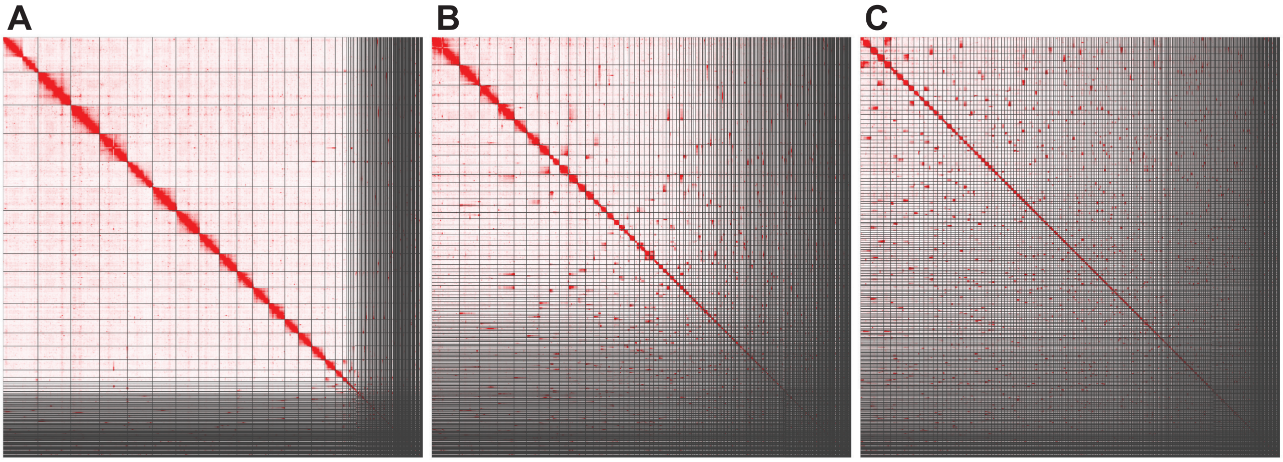


Figure S7: Hi-C contact maps of the humus earthworm (*Lumbricus rubellus*, wcLumRube1) assemblies constructed from YaHS (A), SALSA2 (B) and pin_hic (C).

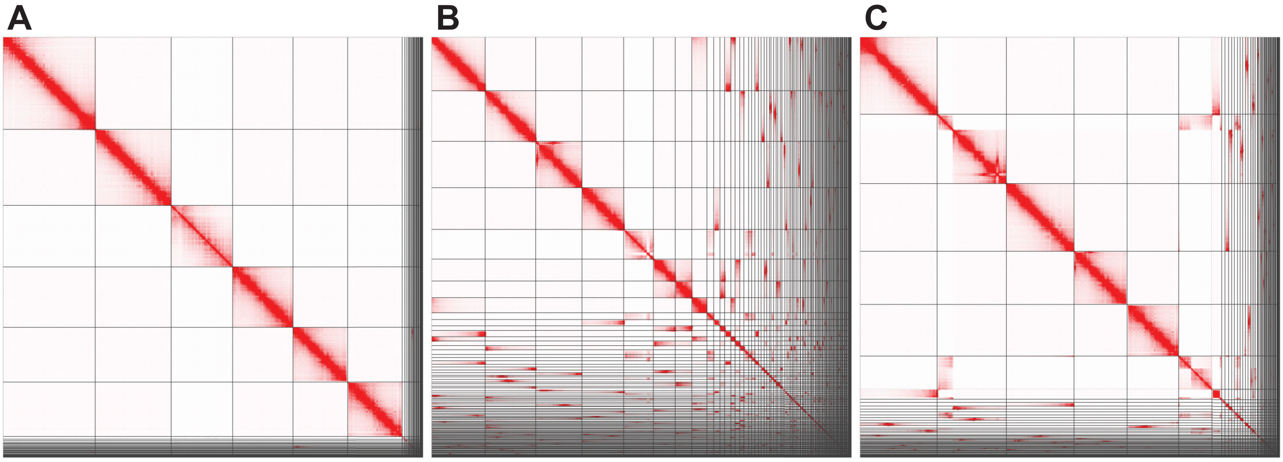


Figure S8: Hi-C contact maps of the hawthorn shieldbug (*Acanthosoma haemorrhoidale*, ihAcaHaem1) assemblies constructed from YaHS (A), SALSA2 (B) and pin_hic (C).

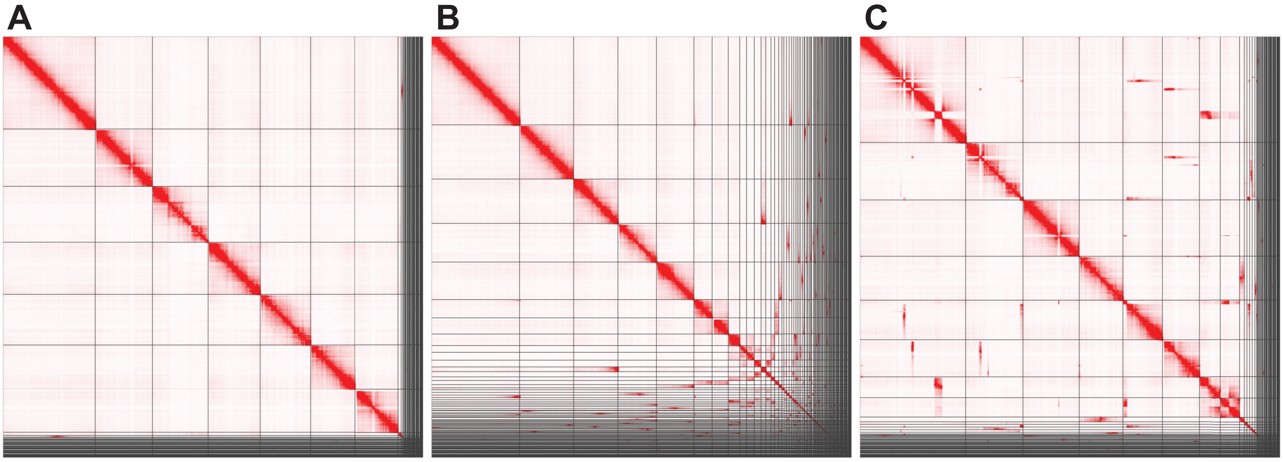


Figure S9: Hi-C contact maps of the *Podabrus alpinus* (icPodAlpi1) assemblies constructed from YaHS (A), SALSA2 (B) and pin_hic (C).

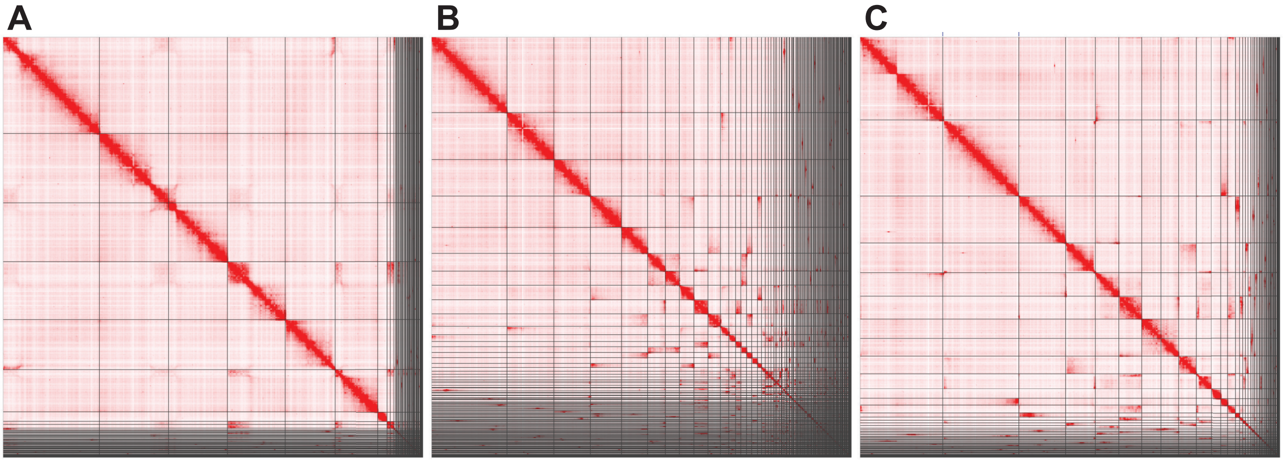


Figure S10: Hi-C contact maps of the dumpy grass hoverfly (*Melanostoma mellinum*, idMelMell2) assemblies constructed from YaHS (A), SALSA2 (B) and pin_hic (C).

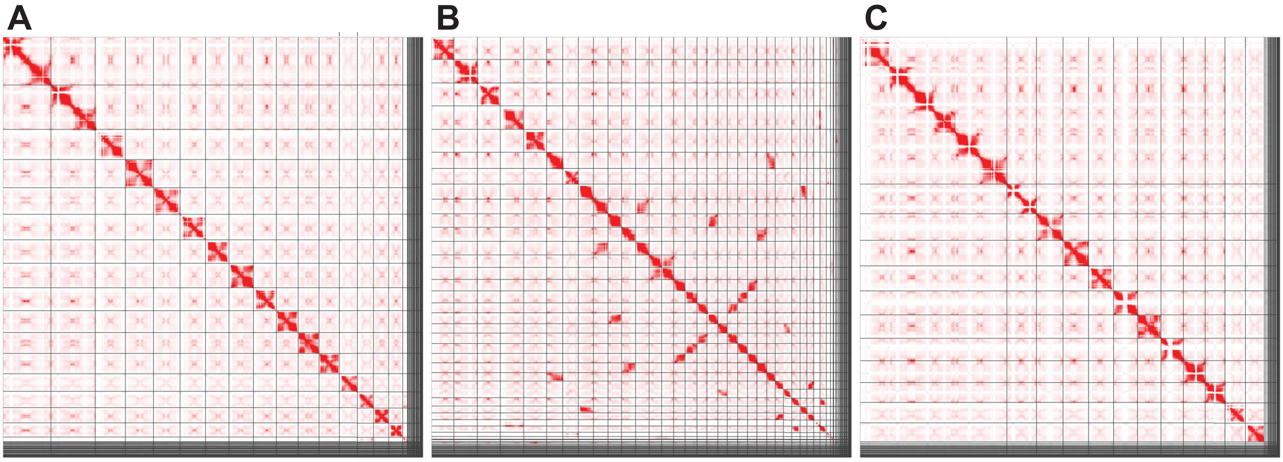


Figure S11: Hi-C contact maps of the *Chamaenerion angustifolium* (drChaAngu1) assemblies constructed from YaHS (A), SALSA2 (B) and pin_hic (C).

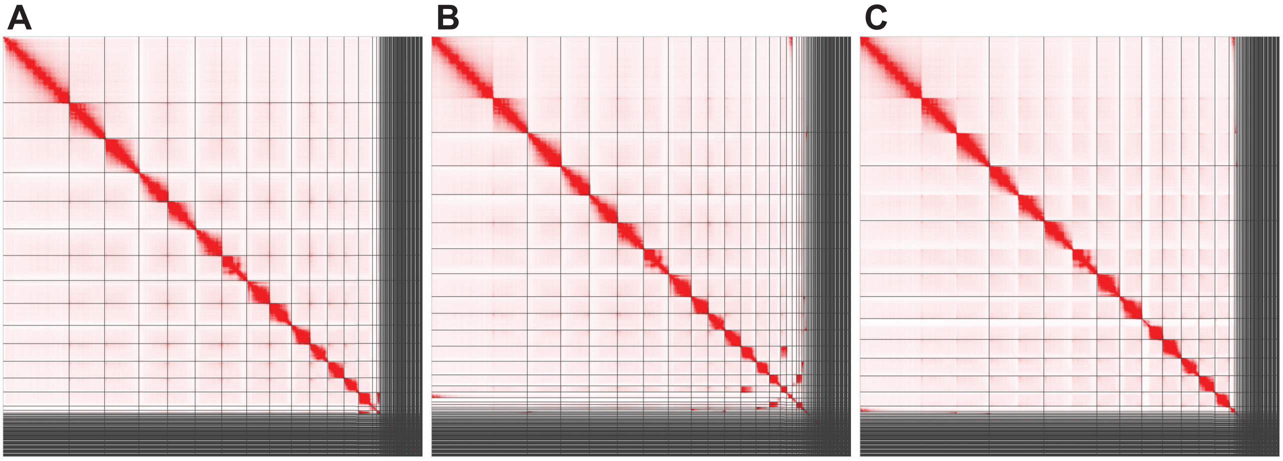


Figure S12: Hi-C contact maps of the ornate tailed digger wasp (*Cerceris rybyensis*, iyCerRyby1) assemblies constructed from YaHS (A), SALSA2 (B) and pin_hic (C).

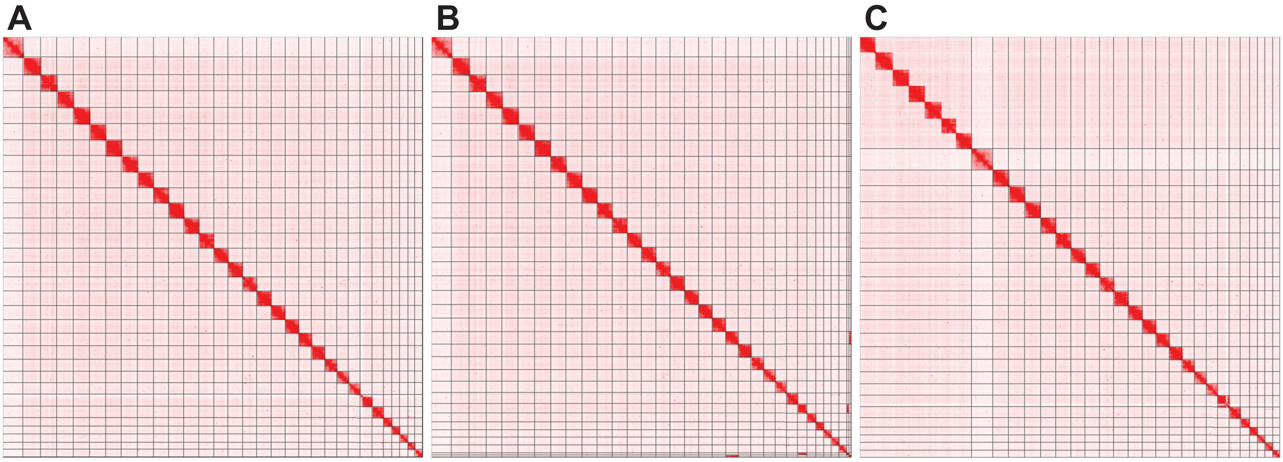


Figure S13: Hi-C contact maps of the diamondback moth (*Plutella xylostella*, ilPluXylo3) assemblies constructed from YaHS (A), SALSA2 (B) and pin_hic (C).

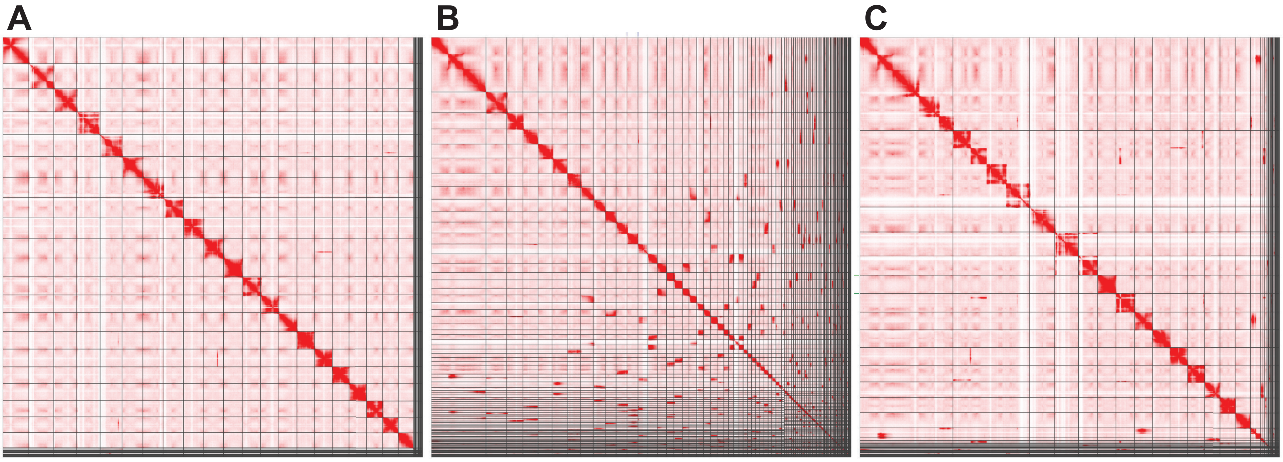


Figure S14: Hi-C contact maps of the soft rush (*Juncus effusus*, lpJunEffu1) assemblies constructed from YaHS (A), SALSA2 (B) and pin_hic (C).

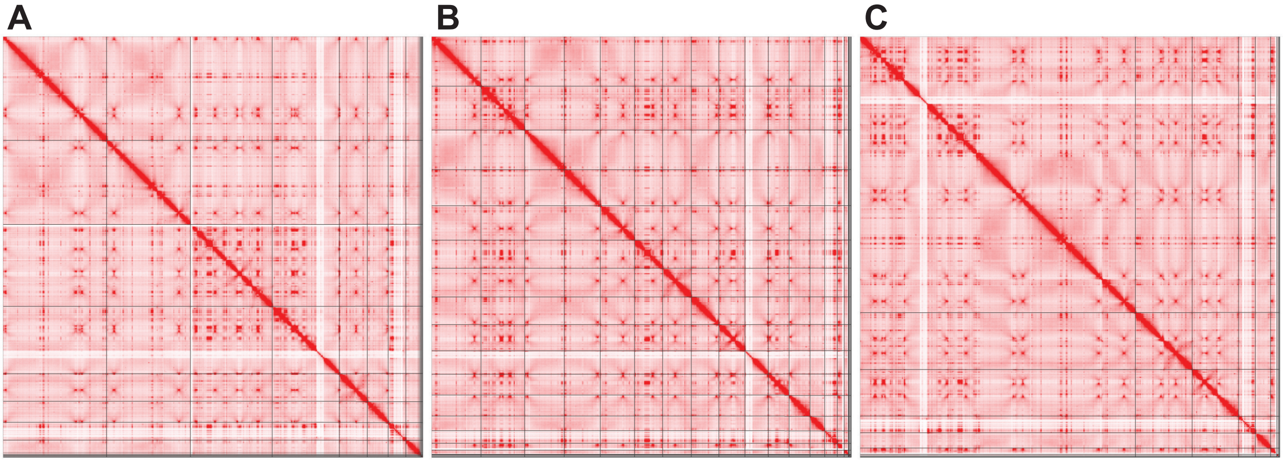


Figure S15: Hi-C contact maps of the chicken of the woods (*Laetiporus sulphureus*, gfLaeSulp1) assemblies constructed from YaHS (A), SALSA2 (B) and pin_hic (C).
